# Conservation and divergence of sex-biased gene expression across 50 million years of *Drosophila* evolution

**DOI:** 10.1101/2025.10.10.681669

**Authors:** Amanda Glaser-Schmitt, John Parsch

## Abstract

In sexually dimorphic species, sex-biased gene expression plays an important role in driving morphological, physiological, and behavioral differences between males and females. Here, we examined patterns of sex-biased gene expression within and among 6 *Drosophila* species with divergence times ranging from 2–50 million years. We detected contrasting patterns of sex bias conservation and turnover between heads and bodies, with more extensive sex-biased expression and greater conservation of sex-biased expression in the body, but more species-specific turnover of sex-biased expression in the head, where conserved, unbiased expression was common. Interestingly, lineage-specific gains of sex-biased expression occurred most often via concordant expression changes in both sexes that differed only in their magnitude, with this pattern being particularly strong in the head, suggesting that the majority of lineage-specific sex bias gains do not represent a resolution of sexual antagonism, but instead reflect regulatory changes shared between the sexes. We detected an enrichment of putatively positively selected expression changes among sex-biased genes in both body parts. Altogether, our findings suggest that sex-biased expression changes are often facilitated by selection, including selection acting on the sex with lower expression, which is especially common for male expression of female-biased genes. We also detected differences in the proportion of sex-biased genes located on the X chromosome depending on sex bias and body part. Overall, our results provide novel insights into the dynamics of sex-biased gene expression, as well as the molecular mechanisms and selective forces underlying its turnover, across short and long evolutionary timescales.

## Introduction

Gene expression variation underlies much of the phenotypic variation that we observe among species, populations, and individuals of the same species (King and Wilson 1975; Wray et al. 2003; Buchberger et al. 2019). In species with sexual dimorphism, this gene expression variation is particularly important as the vast majority or, in some cases, all of the genome is shared by both sexes (Ellegren and Parsch 2007). Therefore, sex-biased (SB) gene expression, i.e. the differential expression of a shared genome between males and females, plays an important role in driving morphological, physiological, and behavioral differences between the sexes (reviewed in Parsch and Ellegren 2013; Grath and Parsch 2016; Tosto et al. 2023). Indeed, quantifying and understanding patterns of sex-biased gene expression over both short and longer evolutionary timescales can help us better understand sexual conflict and the evolutionary forces that shape and maintain phenotypic variation, especially between the sexes, which in turn can shed light on biodiversity and the evolution of complex phenotypic traits (reviewed in Tosto et al. 2023). For example, sex-specific gene expression and regulation have been implicated in the susceptibility and presentation of several human diseases (e.g. Shi et al. 2016; Khodursky et al. 2022; Fass et al. 2024), as well as variation in human height (Naqvi et al. 2019), all of which represent complex, highly polygenic phenotypes.

Studies have shown that sex-biased gene expression is extensive (Meiklejohn et al. 2003; Yang et al. 2006; Oliva et al. 2020), can be tissue-, developmental stage-, and cell type-specific (Yang et al. 2006; Ma et al. 2018; Naqvi et al. 2019; Oliva et al. 2020; Khodursky et al. 2022; Rodríguez-Montes et al. 2023), and evolves rapidly (Assis et al. 2012; Whittle and Johannesson 2013; Harrison et al. 2015; Yang et al. 2016; Xie et al. 2025). The shared genome has long been thought to impose genetic constraints on the sexes (Lande 1980; Griffen et al. 2013), and SB gene expression is thought to be an indication of this ongoing sexual conflict (i.e. sexual antagonism) or its resolution (Connallon and Knowles 2005; Innocenti and Morrow 2010). Indeed, previous studies of SB gene expression in diverse taxa have found that sexual conflict is often resolved via SB gene expression and/or sex-specific regulation (e.g. Cheng and Kirkpatrick 2016; Dean et al. 2016; Wright et al. 2018).

The model insect, *Drosophila melanogaster*, was utilized in some of the earliest studies of SB gene expression (Jin et al. 2001; Arbeitman et al. 2002; Ranz et al. 2003; Parisi et al. 2003) and has continued to be an important model system for studying its evolution at both the interspecific and intraspecific levels (e.g. Meiklejohn et al. 2003; Ranz et al. 2004; Gibson et al. 2004; Catalán et al. 2012; Llopart et al. 2012; Meiklejohn et al. 2014; Huylmans and Parsch 2014; Khodursky et al. 2020). Despite this progress, transcriptomic studies that systematically compare SB gene expression among species while also assessing the full range of within-species expression diversity are currently lacking. Previous studies in the genus *Drosophila* were restricted in their phylogenetic scale (e.g. Ranz et al. 2003; Grath and Parsch 2012; Müller et al. 2012; Khodursky et al. 2020) or had limited samples for determining within-species expression variation (e.g. Zhang et al. 2007; Pal et al. 2023).

The characterization of expression variation both within and between species is important for inferring the type of selection that influences gene expression, as the 2 levels of variation are thought to represent, respectively, the short-term and long-term outcomes of the same underlying evolutionary process (Kimura 1983; Hudson et al. 1987; McDonald and Kreitman 1991). For this reason, we measured gene expression in 2 body parts (head and body) of adult males and females of 6 *Drosophila* species spanning divergence times of up to 50 million years (Tamura et al. 2004). Within each species, we included 5–8 strains derived from diverse geographic locations. These data allowed us to quantify the conservation and turnover of SB gene expression across the genus in samples that included or excluded the gonad, which is the tissue with the highest proportion of SB genes (Parisi et al. 2004), and address 3 major questions: *i*) How much turnover is there in SB gene expression across the *Drosophila* phylogeny? *ii*) How do genes gain SB expression? For example, does expression increase/decrease in only 1 sex, or does it change in both sexes? If it is the latter, does expression change in opposite directions in the 2 sexes, which is expected if SB expression resolves sexual conflict, or are there concordant expression changes in the 2 sexes, which would be indicative of shared regulatory constraints? *iii*) Do lineage-specific expression changes show molecular signatures of positive selection? If so, does selection occur predominantly in the sex that expression is biased towards? Answering these questions will shed light on the evolutionary mechanisms responsible for the maintenance and turnover of SB gene expression in the *Drosophila* genus.

## Results

In order to characterize SB expression over a range of timescales, we performed RNA-seq of male and female heads and bodies of 6 *Drosophila* species with divergence times ranging between 2–50 million years (Fig 1), including *D. melanogaster* (*Dmel*), *D. simulans* (*Dsim*), *D. suzukii* (*Dsuz*), *D. ananassae* (*Dana*), *D. subobscura* (*Dsub*), and *D. immigrans* (*Dimm*). To get a comprehensive view of expression variation within each species, we sampled 5–8 strains from a broad geographic distribution, encompassing strains from multiple populations across 3 continents, including strains from the putative ancestral range for *Dmel*, *Dsim*, *Dana*, and *Dsuz* (Table S1). We analyzed expression in 9,016 1:1 orthologs present in all species (Thiébaut et al. 2024), 7,570–8,161 of which were expressed in the head and 8,936–9,006 of which were expressed in the body of each species. To get an overview of expression variation, we performed principal component analyses (PCAs) for all samples, within each body part, or within each species. When considering the full dataset, samples clustered strongly by body part (Fig 2A). In the head, the sexes showed only slight separation (Fig 2B), while in the body, males and females were clearly differentiated (Fig 2A, C). Within each sex and body part, strains of each species clustered together, although these clusters often overlapped among species. *Dimm*, which is the most distantly related to the other species (Fig 1), also showed the greatest interspecific differentiation (Fig 2B, C). Within each species, patterns were similar, with body parts and sexes clustering by sample type but differentiation between the sexes much stronger in the body than the head (Fig S1).

**Fig 1:**
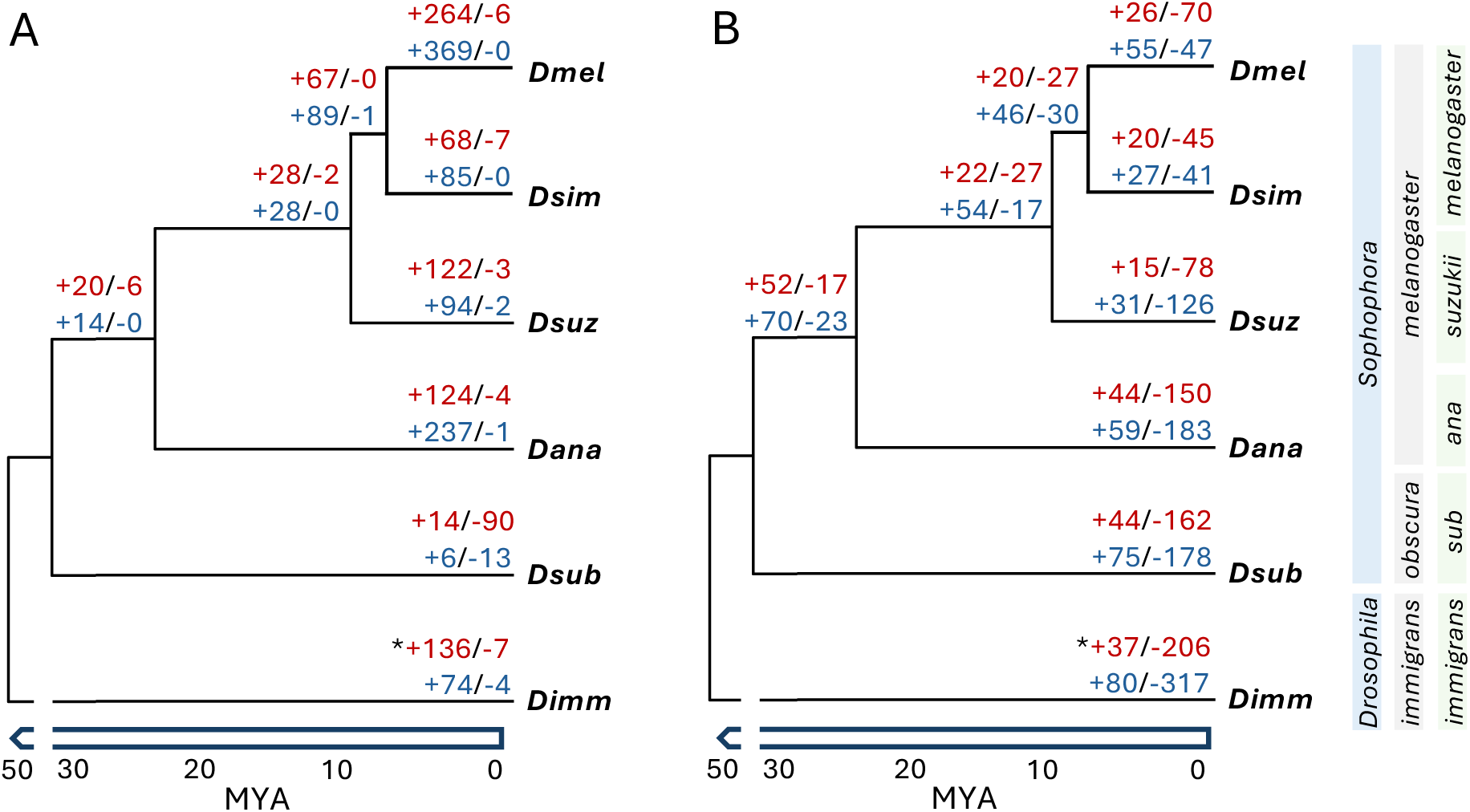
Phylogeny and SB expression turnover in the examined *Drosophila* species in the A) head and B) body. Branch lengths are shown only roughly to scale in millions of years ago (MYA). Numbers at each tip or node indicate sex bias gains (+) and losses (-) for female-(red) and male-biased (blue) genes. The subgenera are shown in the light blue boxes, the species groups in the gray boxes, and the subgroups in the light green boxes (*subobscura*: *sub*; *ananassae*: *ana*). *Gains and losses could not be polarized for *Dimm* due to the lack of an outgroup; therefore, the number of genes with different sex bias relative to the other species is shown.

**Fig 2:**
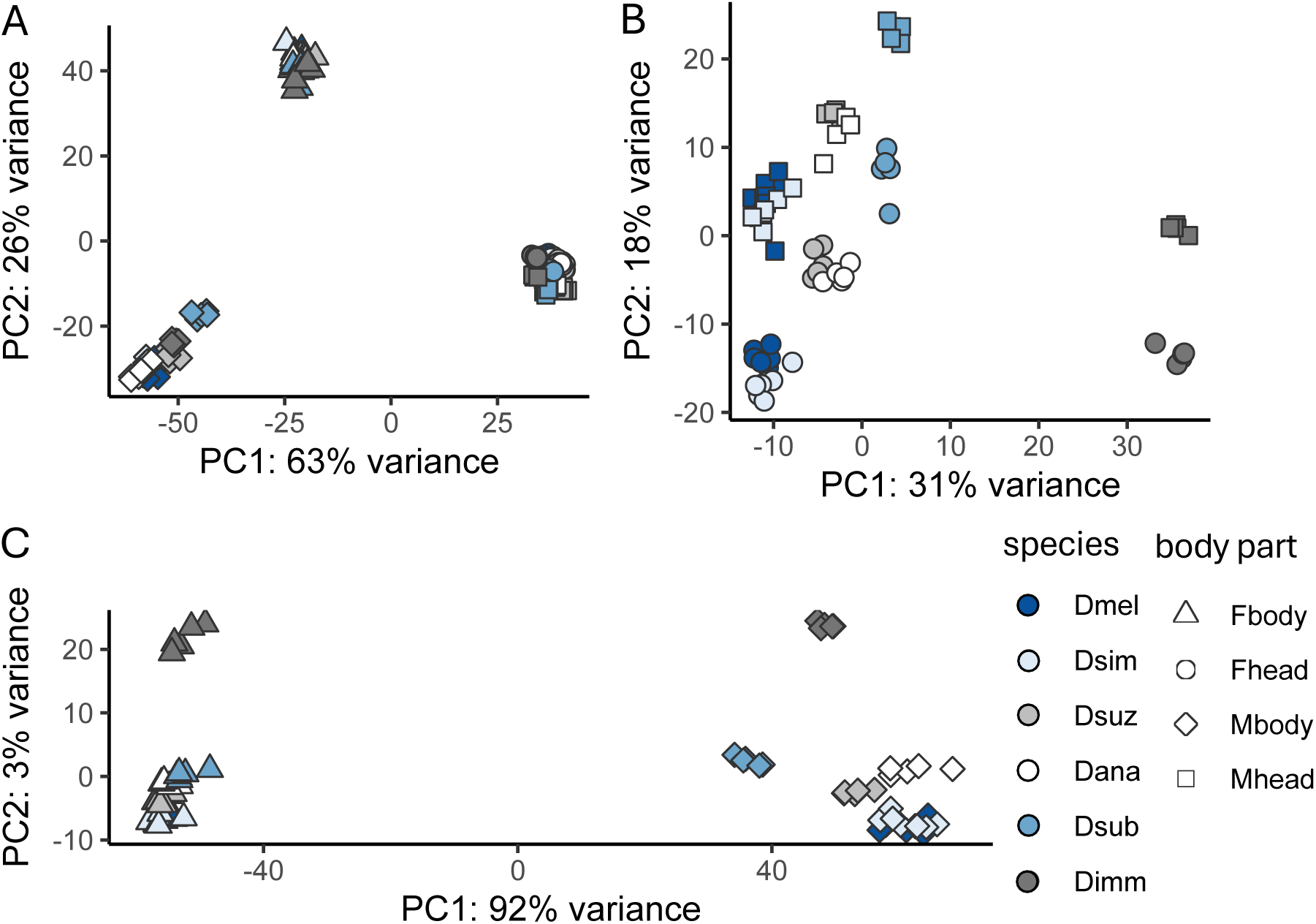
PCA of gene expression profiles in A) all samples, B) the head, and C) the body. The legend at the bottom right indicates the color associated with each species and the shape indicates the sex (male: M; female: F) and body part.

### Patterns and magnitude of sex-biased expression differ among body parts and species

In order to compare overall prevalence, patterns, and degree of sex bias within and among the examined species, we used DESeq2 (Love et al. 2014; see Materials and Methods) to identify male-biased (MB), female-biased (FB), and unbiased (UB) genes within each species. SB gene expression was extensive in the body, with 71.9–81.6% of genes showing sex bias, but less common in the head, with 2.0–21.7% of genes showing sex bias (Table 1). In comparison to the other species, sex bias in *Dsub* heads was very low (2.0% versus 10.1– 21.7%; Table 1), especially when considering MB genes (32 versus 212–814 genes; Table 1). We detected more genes as SB in the *Dmel* head in comparison to the other species, but this excess was most likely driven by the sampling of additional strains in this species rather than any biological differences between species, as down-sampling resulted in similar numbers of SB genes as detected in the other species (Table S2). Because SB gene expression is often tissue-specific (Oliva et al. 2020), we compared sex bias between body parts. Due to the large amount of sex bias in the body, the majority of genes that could be examined in both body parts (64.2–73.9%) were SB in the body but not the head (Fig 3C, Fig S2). The next largest category of overlap was genes UB in both the body and head (14.1–25.2%), with genes showing the same sex bias in both body parts being the third largest category (0.9– 13.0%). Interestingly, genes with opposite sex bias in the head and body were more common than genes SB in the head but not the body (0.7–5.6% versus 0.4–3.1%, respectively). Thus, genes SB in the head were 3–6 times more likely to also be SB in the body, and this sex bias was often, but not always, in the same direction (Fig 3C, Fig S2).

**Fig 3:**
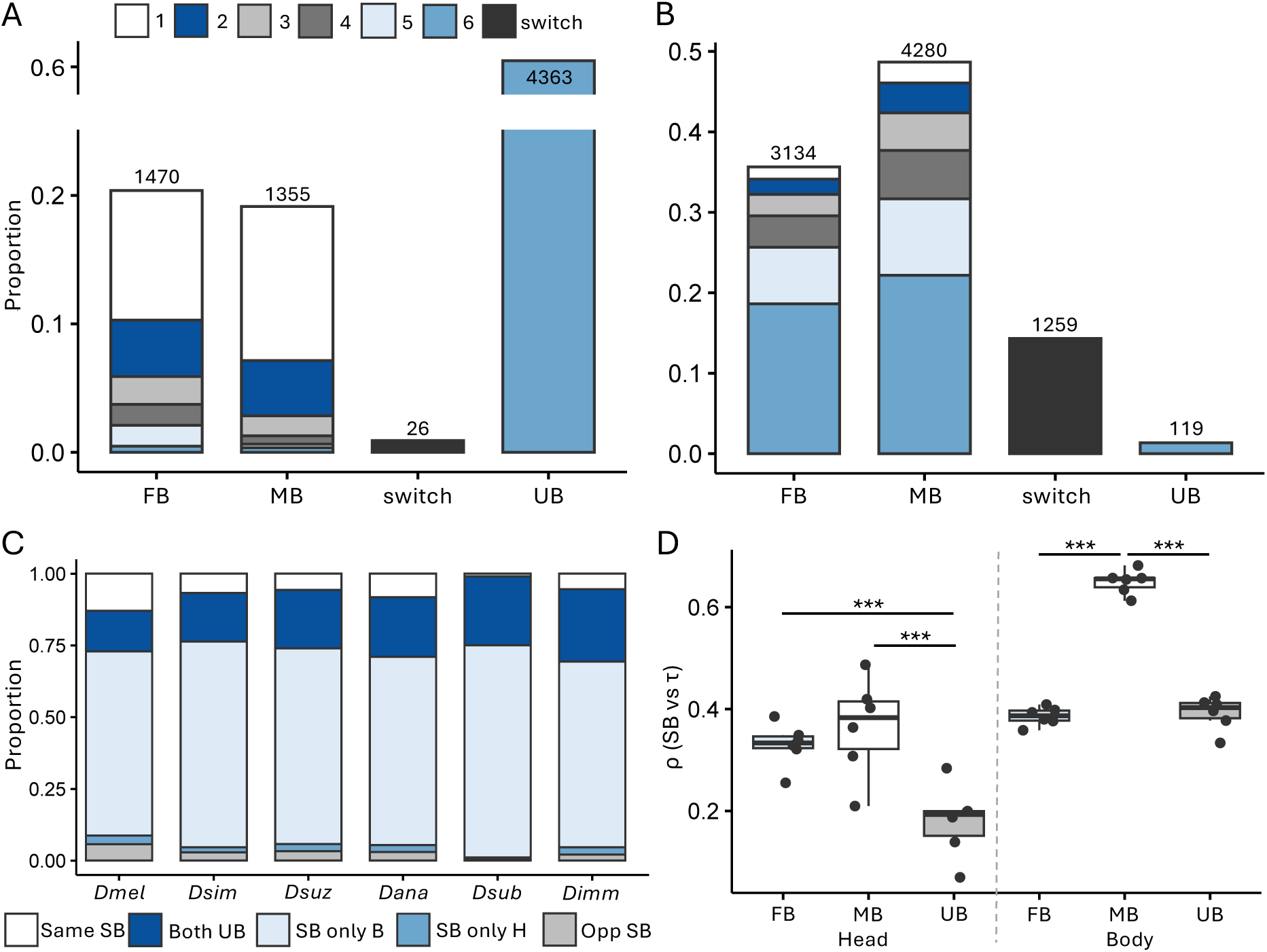
Sex bias within and among species. A,B) Proportion of genes with overlapping SB or UB expression among species in the A) head and B) body. Shown are the proportion of overlapping genes in the same SB category in 1 (white), 2 (dark blue), 3 (light grey), 4 (dark grey), 5 (light blue), and all 6 (medium blue) examined species. Proportion of analyzed genes that switched SB (FB to MB or vice versa) among any of the examined species is shown in black. The number of total genes in each category are shown above the bar. C) Proportion of overlapping genes that had shared SB (same SB; white) or UB (both UB; dark blue) expression, were SB in only the body (SB only B; light blue) or head (SB only H; medium blue), or opposite sex bias between the head and body (opp SB; light grey) are shown. D) Spearman’s ρ correlations between τ and magnitude of sex bias in the head and body for MB, FB, and UB genes. Significance was assessed with a Mann-Whitney *U* test. \*\*\**P* < 0.005. Non-significant comparisons not shown.

**Table 1:**
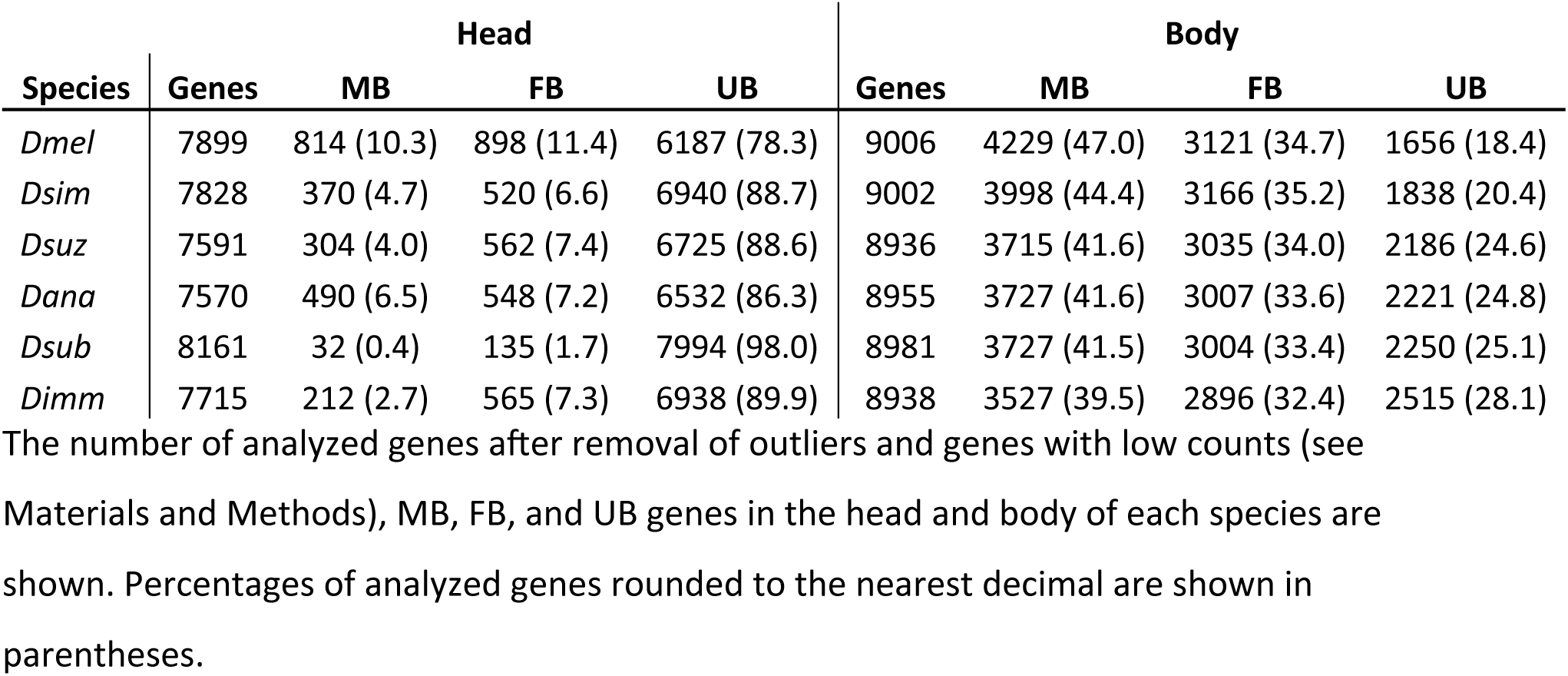
Sex bias in the head and body of 6 *Drosophila* species.

To determine if the magnitude of sex bias differed among body parts, the sex in which sex bias was detected, or species, we compared the degree of sex bias (i.e. the absolute value of our sex bias estimates) among categories. The magnitude of sex bias was significantly higher in the body than the head for all species for both MB and FB genes (*t*-test; BH corrected *P* < 1.5 × 10^−15^ for all; Fig S3). In the head, the degree of sex bias was higher for FB than for MB genes, while in the body, the opposite pattern was detected, which was significant for all comparisons in all species (*t*-test; BH corrected *P* < 4.5 × 10^−7^ for all; Fig S3), except *Dana* and *Dsub* heads (BH corrected *P* = 1). We also detected significant differences in the degree of sex bias within the same body part and sex bias category (MB or FB) among species, especially in the head, for many but not all comparisons (Fig S3). To determine how these differences in the magnitude of sex bias relate to differences in overall expression, we calculated the overall expression divergence (as measured by 1 – ρ) between body parts and species. Divergence between sexes was lowest in the head within species (0.8–1.3%; Table S3), while divergence in the body was fairly similar to divergence between body parts regardless of sex (27.7–36.5% versus 25.2–43.6%%; Tables S3–S4). Divergence between species within the same body part and sex was higher than in the head but lower than in the body between sexes within species (1.8–12.0%; Tables S3–S4). Thus, we detected differences in the magnitude of expression bias based on the type of sex bias, the body part, and the species, but overall expression levels showed the strongest correlation (between sexes in the head) and weakest correlation (between the examined body parts in either sex) within species.

### Tissue specificity and expression level are associated with differences in sex bias

Previous studies have shown that SB genes often show a high degree of tissue specificity (Meisel 2011; Dean et al. 2016; Khodursky et al. 2020). We therefore used the adult *Dmel* tissue expression data in FlyAtlas2 (Krause et al. 2022) to examine expression of our SB genes in individual tissues (Table S5). As might be expected for SB genes, MB genes in the head or body were often most highly expressed in testes or a somatic tissue, while FB genes were most highly expressed in the ovary or a somatic tissue. To better understand the relationship between expression level, breadth, and sex bias in our dataset, we first examined the overall expression level of SB and UB genes in the head and body of each species. We detected significant effects of both species and sex bias category on the expression level in both body parts (ANOVA; *P* < 1.5 × 10^−6^ for all; Fig S4). In both the head and body, FB genes were the most highly expressed, followed by MB genes, and then UB genes, except in *Dsub* heads, where MB genes were the most lowly expressed (Fig S4). Thus, differences in sex bias were associated with differences in overall gene expression levels.

Next, to better understand the association between sex bias in our dataset and tissue-specificity, we used the FlyAtlas2 adult tissue expression data (Krause et al. 2022) to calculate the tissue specificity index, τ, which ranges from 0 to 1 with values closer to 1 indicating higher tissue specificity, for each gene. To determine if genes we identified as SB were more tissue-specific than UB genes as previous studies have found (Meisel 2011; Dean et al. 2016; Khodursky et al. 2020), we compared tissue specificity among SB and UB genes in each species and body part. We detected a significant effect of sex bias category in both body parts (ANOVA; *P* < 1.6 × 10^−10^ for all; Fig S5), suggesting that genes in different sex bias categories display different levels of tissue specificity. Tissue specificity was highest for MB genes in both the head and body for all species (Fig S5). In the head, τ was mostly similar between FB and UB genes, while in the body, τ was lower in FB than in UB genes (Fig S5). Next, we examined if the magnitude of sex bias was associated with the degree of tissue specificity as well as the sex bias category. The magnitude of sex bias was significantly positively correlated with τ in both body parts in all sex bias categories and all examined species (Spearman’s ρ; *P* < 0.003 for all) except MB genes in the head of *Dsub* (*P* = 0.2483). Across all species, this correlation between the degree of sex bias and τ was highest for MB genes in both body parts (Fig 3D), which was significantly higher than UB genes in both body parts and FB genes in the body (Mann-Whitney *U* test; *P* < 0.005 for all), while the correlation for FB genes was only significantly higher than UB genes in the head (*P* < 0.005; Fig 3). Thus, tissue specificity was positively associated with sex bias, with the strongest correlation in MB genes, which were also the most tissue-specific. This correlation was particularly strong in the body, which might be driven at least in part by the presence of the testes, which is the tissue with the highest proportion of MB genes (Parisi et al. 2004).

### Patterns of sex bias turnover among species differ between head and body

To better understand how sex bias is conserved among species, we identified genes that were SB or UB in all 6 species. In the head, we identified 4,363 (53.5–57.6%) genes with conserved UB expression in all species, but only 1 (< 0.02%) and 35 (< 0.5%) genes as having conserved MB or FB expression, respectively. On the other hand, we identified only 119 (1.3%) genes with UB expression in all species in the body, but 1,638 (18.2–18.3%) and 1,949 (21.6–21.8%) genes with conserved FB and MB expression, respectively. Next, to understand how turnover in sex bias occurs among species, we employed a conservative approach (see Materials and Methods) to identify lineage-specific gains, losses, or switching of FB or MB expression that occurred on only 1 branch of the phylogeny (i.e. changes that occurred only at 1 internal or external node of the phylogeny; Fig 1, Table S6). In the head, we detected many more sex bias gains than losses for all species except *Dsub* (Fig 1A; 1,629 total lineage-specific gains versus 135 losses, 103 of which were in *Dsub*); while, in the body we detected more sex bias losses than gains (1,227 losses versus 663 gains; Fig 1B), although the pattern was not as extreme as in the head. Across the entire phylogeny, only 171 genes switched sex bias on a specific lineage, all of which occurred in the body and the majority of which (64.3%) switched from FB to MB (Table S6).

To better understand how the sex bias of a gene differs among species across the entire phylogeny, we examined the of genes analyzed in all species that were SB or UB in 1–6 of the examined species. In the head, genes that were SB in 1 species were often UB in the other species (10.1–12.0% of all analyzed genes; Fig 3A), but if a gene was SB in more than 1 species, it almost always shared the same sex bias between species (0.01–12.0% of genes depending on the number of species in which a gene was detected as SB, 6.8–10.3% in total; Fig 3A). On the other hand, only 26 genes switched their sex bias (i.e. from MB to FB or vice versa) between species (0.4% of all analyzed genes; Fig 3A). In the body, SB genes were often shared between species, with the majority showing sex bias towards the same sex in at least 2 species (1.5–22.2% of genes depending on the number of species in which a gene was detected as SB, 34.1–46.1% in total; Fig 3B). A larger proportion of genes switched sex bias in the body (14.3% of all analyzed genes; Fig 3B) compared to the head. To determine how widespread this switching was among individual species, we performed pairwise comparisons of sex bias in all examined genes between species for each body part (Figs S7, S8). A small proportion (1.4–4.5%; Fig S8) of analyzed genes switched SB, suggesting that relatively few genes switch sex bias among the individual species, but this number adds up across the phylogeny. However, we should note that because the body is a composite of multiple tissues, it is possible that some of these switches represent opposing changes in different tissues rather than a switch in the same tissue. Overall, we detected a relatively large amount of sex bias turnover in both body parts among species, which occurred in a mostly species-specific manner in the head (Figs 1, 3). Moreover, when turnover occurred in the head, a gene’s expression most often switched between being UB and biased towards a single sex, but rarely between the 2 sexes.

### Sex bias is most often gained or lost via correlated expression changes in both sexes, especially in the head

In order to better understand how expression changes during the gain or loss of sex bias as well as how the magnitude of an expression change affects these patterns, we examined how expression in a focal species changed relative to the inferred ancestral expression within each sex (see Materials and Methods) for species-specific gains or losses of SB expression (i.e. how do male and female expression levels change when sex bias is gained or lost). In the head, the majority of expression changes occurred in the same direction (either up-regulation or down-regulation) in both sexes (75.6% of all sex bias gains and losses; Fig 4), and this pattern was even stronger for genes with large expression changes (log_2_ fold-change ≥ 1), 95.2% of which had concordant expression changes between the sexes (versus 74.3% of genes with small expression changes; Fig S8). Although concordant expression changes were also more frequent than opposing changes in the body (54.3% versus 45.7%), the direction of expression changes in males versus females was more evenly distributed compared to the head (*P* < 2.2 × 10_-16_; χ^2^ test; Fig 4). Genes with large expression changes (log_2_ fold-change ≥ 2) were more similar to patterns in the head, with 66.7% of genes showing concordant expression changes between sexes (versus 53.3% of genes with small expression changes; Fig S8). If gene expression changes are concordant, we would expect there to be a positive correlation between the sexes. Consistant with this expectation, expression changes associated with SB gains were significantly positively correlated in the head and body in all species (Spearman’s ρ; *P* < 0.05 for all) except for MB gains in the *Dsub* head and FB gains in the *Dmel* and *Dsim* body (Figs S9–S10). This correlation in gene expression changes between the sexes was significantly higher in the head than in the body (paired Mann-Whitney *U* test; *P =* 0.0003). However, this pattern was less apparent for SB losses, for which this correlation was sometimes negative and only significant in half of the comparisons (Figs S9–S10), although in the head this might be at least partially explained by the low number of SB losses in each species. Indeed, correlations in gene expression changes between the sexes were significantly higher for SB gains than for losses in both the head and the body (*P* < 0.05 for all; Fig 4).

**Fig 4:**
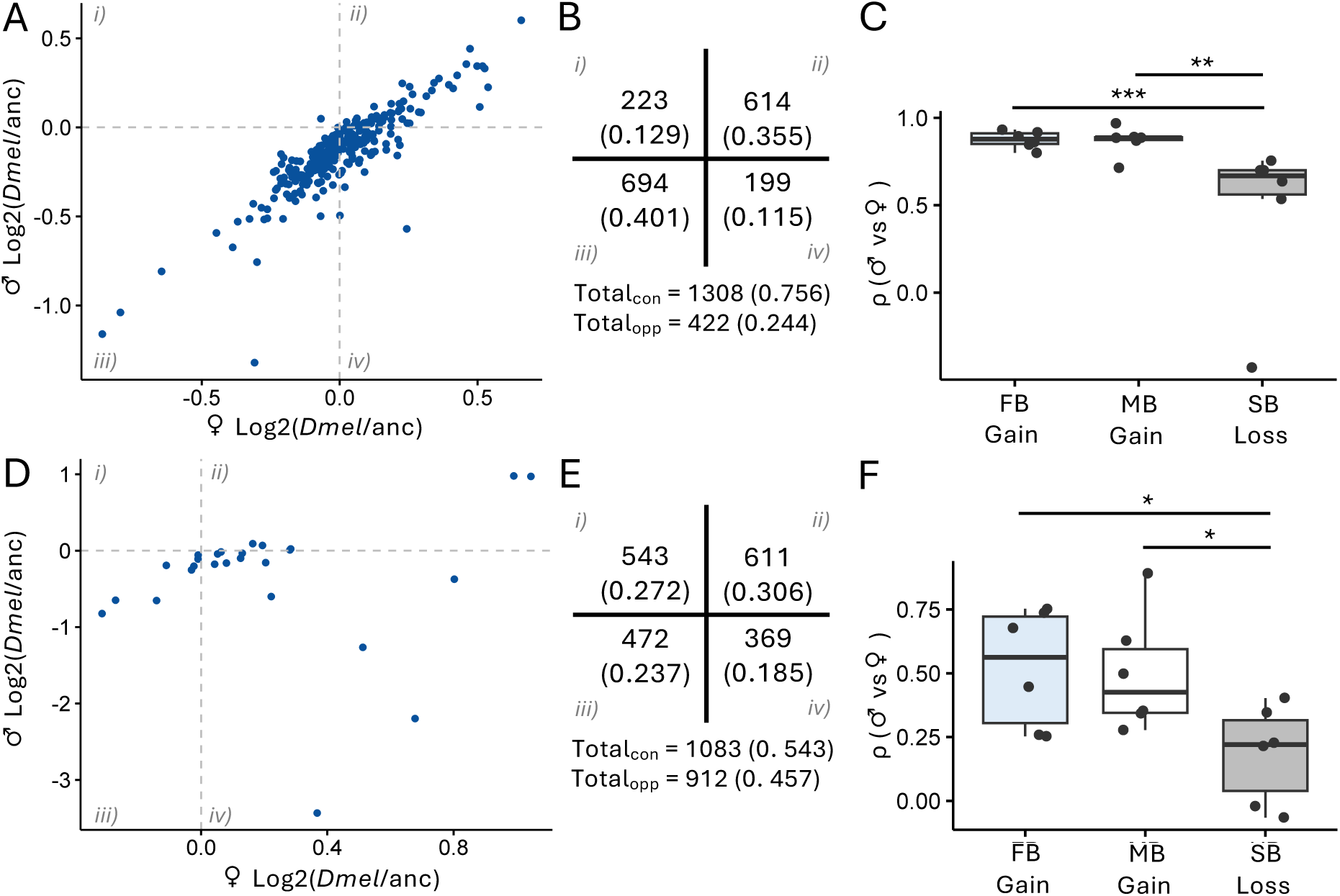
Expression changes underlying species-specific gains and losses of SB gene expression. As an example, scatterplots depicting the change between *Dmel* and ancestral expression in males versus females for genes with FB gains in the A) head and D) body are shown. All other scatterplots can be found in Figs S9–S10. A,B,D,E) Genes with concordant (con) changes in expression between the sexes are located in quadrants ii and iii; while genes with opposing (opp) expression changes between the sexes are located in quadrants i and iv. B, E) Total genes in each quadrant for the entire phylogeny in the B) head and E) body are shown. In parentheses, the proportion of total gene expression changes are shown. C, F) Spearman’s ρ correlations between male and female expression changes within each species for SB gains and losses in the C) head and F) body. Significance was assessed with a Mann-Whitney *U* test. \*\*\**P* < 0.005, \*\**P* < 0.01, \**P* < 0.05. Non-significant comparisons not shown.

Next, we examined how the size of expression changes and ancestral sex bias affect the distribution of expression changes during SB loss or gain. In other words, are large expression changes more likely to be driven by up- or down-regulation of gene expression and how does ancestral sex bias affect these patterns? For the majority of large gene expression changes (log_2_ fold-change ≥ 1 in the head or ≥ 2 in the body; 56.7–75.2% of genes), gain or loss of sex bias occurred via a down-regulation in both sexes, with only 10– 20% of genes up-regulated (Fig S8). In contrast, up- or down-regulation in both sexes was more evenly distributed for genes with small expression changes (32.3–36.5% and 21.0– 37.8% of genes for head and body, respectively; Fig S8). Due to the lack of SB reversals and the low number of SB losses for most species in the head, it was difficult to determine if the ancestral sex bias of a gene affected the distribution of expression changes, although MB and FB losses tended to overlap with no clear separation based on the ancestral sex bias (Fig S9). We observed a similar pattern for MB and FB gains in the body, with no clear clustering based on whether sex bias was gained from an UB or SB gene (Fig S10). On the other hand, FB and MB losses in the body were clearly separated (Fig S10), with genes that lost female bias tending to show down-regulation in females and/or up-regulation in males and the opposite for MB losses. Thus, a gain or loss of sex bias most often involved an expression change in the same direction (up-regulation or down-regulation) in both sexes, but of greater magnitude in 1 sex than the other, with larger expression changes tending to be associated with down-regulated expression in both sexes. Overall, reversal-of-expression changes were less common but were almost as frequent as concordant expression changes in the body.

### Contrasting distributions of X versus autosomal SB genes in the head and body

In male heterogametic species, such as *Drosophila*, females have 2 copies of the X chromosome while males have only 1, which can lead to distinctive patterns of gene expression and content on the X chromosome versus the autosomes. Indeed, previous studies found that, in whole flies and gonads, FB genes were overrepresented on the X chromosome, while MB genes were underrepresented (Parisi et al. 2003; Ranz et al. 2003; Sturgill et al. 2007). In contrast, in the head and brain, genes with SB expression, and especially MB expression, were enriched on the X chromosome (Catalán et al. 2012; Huylmans and Parsch 2015; Khodursky et al. 2020). We detected similar patterns across the species included in our study, with the proportion of X-linked genes depending upon the body part and, in some cases, the species in which SB gene expression was detected (Fig 5). In the head, SB genes were generally over-represented on the X chromosome, but whether this enrichment occurred for MB or FB genes depended upon the species, and in some cases (e.g. FB genes in *Dsuz* or MB genes in *Dana*) there was a slight underrepresentation of SB genes on the X (Fig 5A). Interestingly, in *Dsub* MB genes were highly enriched on the X chromosome but highly underrepresented on the autosomes (Fig 5A). In the body, FB genes were enriched, while MB genes were underrepresented on the X chromosome for all species (Fig 5B). Reciprocally, MB genes were enriched while FB genes were underrepresented on the autosomes for all species except *Dana* (Fig 5B). Thus, SB genes displayed unique chromosomal distributions depending on the examined body part.

**Fig 5:**
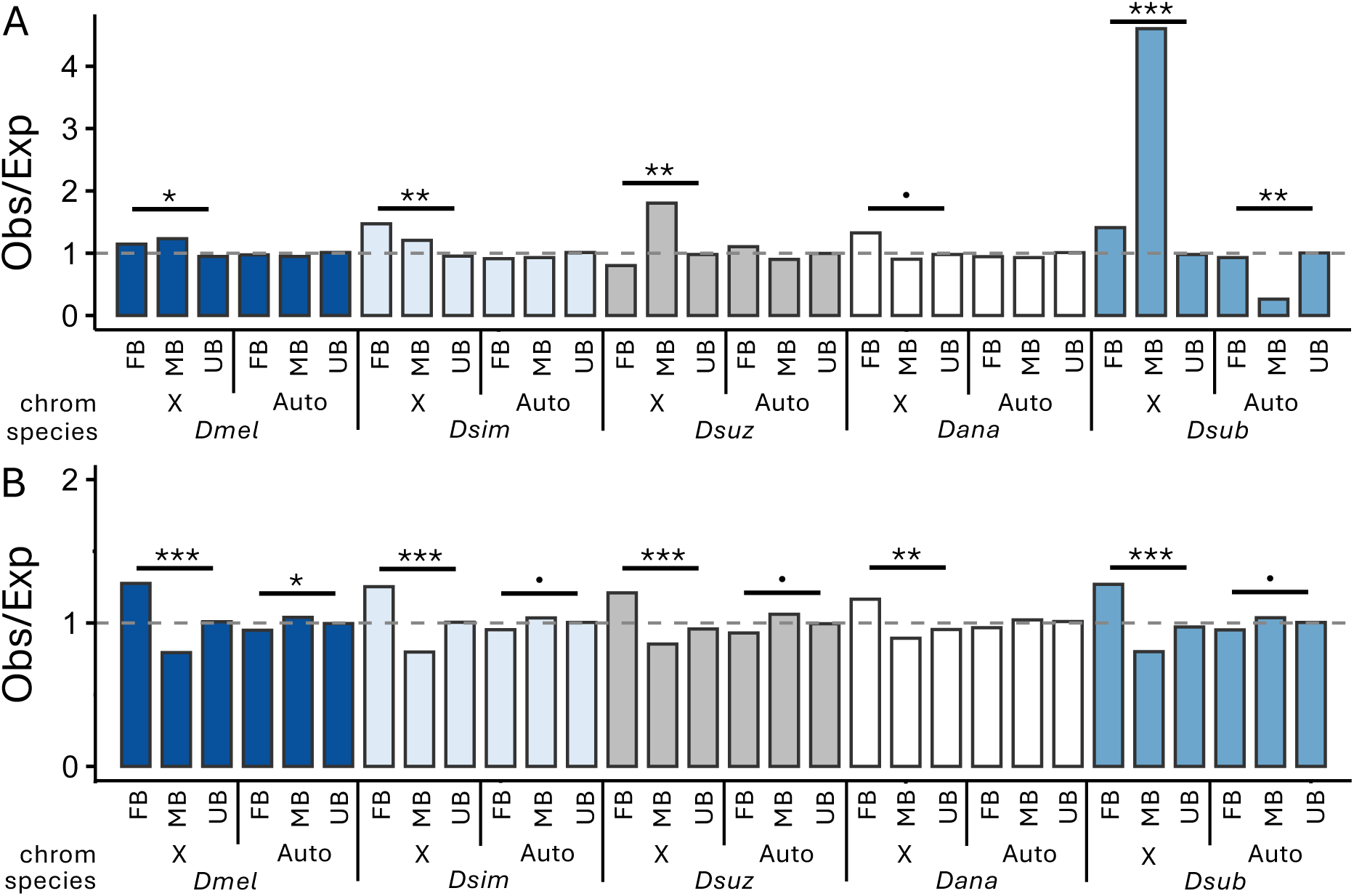
Chromosomal distribution of SB genes in the A) head and B) body. Shown are the ratios of observed (obs) to expected (exp) number of genes in each SB category on the X versus the autosomes based on sex bias category and the number of analyzed on the X versus the autosomes. Significance was assessed with a χ^2^ test. \*\*\**P* < 0.005, \*\**P* < 0.01, \**P* < 0.05, marginally non-significant: .*P* < 0.1. Non-significant comparisons not shown.

### Signs of positive selection at the gene expression level are pervasive in SB genes

A previous study in the brain of 3 *melanogaster* subgroup species found that SB genes, and MB genes in particular, are enriched for signatures of directional selection (Khodursky et al. 2020). In order to test for signs of putative positive selection at the gene expression level in our dataset, we calculated Δx (see Materials and Methods) in both sexes for all species relative to the outgroup species *Dimm*. Briefly, Δx is a measure of the ratio of expression divergence to polymorphism within a species. An absolute Δx value greater than 1 indicates that divergence of expression in the focal species from *Dimm* exceeds expression polymorphism within that species, indicating that positive selection may have occurred, with Δx < −1 representing a down-regulation in the focal species and Δx > 1 indicating an up-regulation. This measure has previously been applied to detect directional selection at the gene expression level in multiple diverse taxa (e.g. Moghadam et al. 2012; Zemp et al. 2016; Lichilín et al. 2021; Cossard et al. 2022).

We detected putative signs of positive selection (Δx > 1 or < −1) in 6.3–76.5% of genes depending upon the sex, species, and sex bias category (Table S7; Fig S11), with the proportion of putatively positively selected genes higher for both FB and MB than for UB genes for all comparisons. Indeed, we detected significantly more FB and/or MB genes as putatively positively selected than expected by chance in both sexes in the head for all species (Fig 6A; χ^2^ test; *P* < 0.05), except in *Dana* females and *Dsub* males, which were marginally non-significant (*P* < 0.09). Putatively positively selected MB or FB genes were similarly enriched to a lesser extent in 1 or both sexes in all species in the body, although we detected fewer FB genes as putatively positively selected than expected in *Dsuz* females (Fig 6B). This enrichment of putatively positively selected MB and/or FB genes occurred on both the X and the autosomes, although in some cases we also detected fewer SB genes than expected both on the X and the autosomes, especially in the body (Fig S12). Overall, this slightly higher enrichment of selected SB genes in the head suggests that although we detected less SB expression in head, the genes that did show SB expression were more likely to experience positive selection. Interestingly, SB genes were often enriched for putative positive selection in the sex in which they were detected as SB (Fig 6A, B), but not always. Unexpectedly, SB genes with signs of positive selection were often also (and sometimes only) enriched in the opposite sex (i.e. the sex with lower expression; Table S8). Indeed, they were enriched in at least 1 sex in all species in both the head and body, although this enrichment was marginally non-significant in females in the *Dsub* body (Fig 6A, B, Table S8). To assess differences in selection between the sexes, we compared Δx between males and females. We detected significant differences in the magnitude of Δx for the majority of sex bias categories and species in both head and body (80% for both; Fig 6C, D), with the absolute value of Δx most often shifted higher in males, especially in the body (75 and 92% of significant shifts in the head and body, respectively), and the mean absolute value in the significantly higher sex usually close to or above 1 (88% of significant shifts; Fig 6C, D; Fig S11). Despite this shift in magnitude, Δx was significantly positively correlated between the sexes in all sex bias categories and species (Spearman’s ρ; *P* <2.2 × 10^−16^ for all; Table S8), except *Dsub* MB genes in the head (*P* = 0.153), suggesting selection in the 2 sexes is positively associated and, in general, selection drives gene expression in the same direction in both sexes. Moreover, these positive correlations were significantly stronger in the body than in the head (mean of 0.827 versus 0.585 in the body and head, respectively; paired Mann-Whitney U test; *P* = 5.37 × 10^−3^). Thus, we detected an enrichment of putatively positively selected genes among SB genes in both the head and body, and this enrichment often occurred in either both sexes or only the sex with lower expression, mainly involving the male expression of FB genes. Moreover, the overall magnitude of Δx was significantly higher in males in comparison to females, especially in the body.

**Fig 6:**
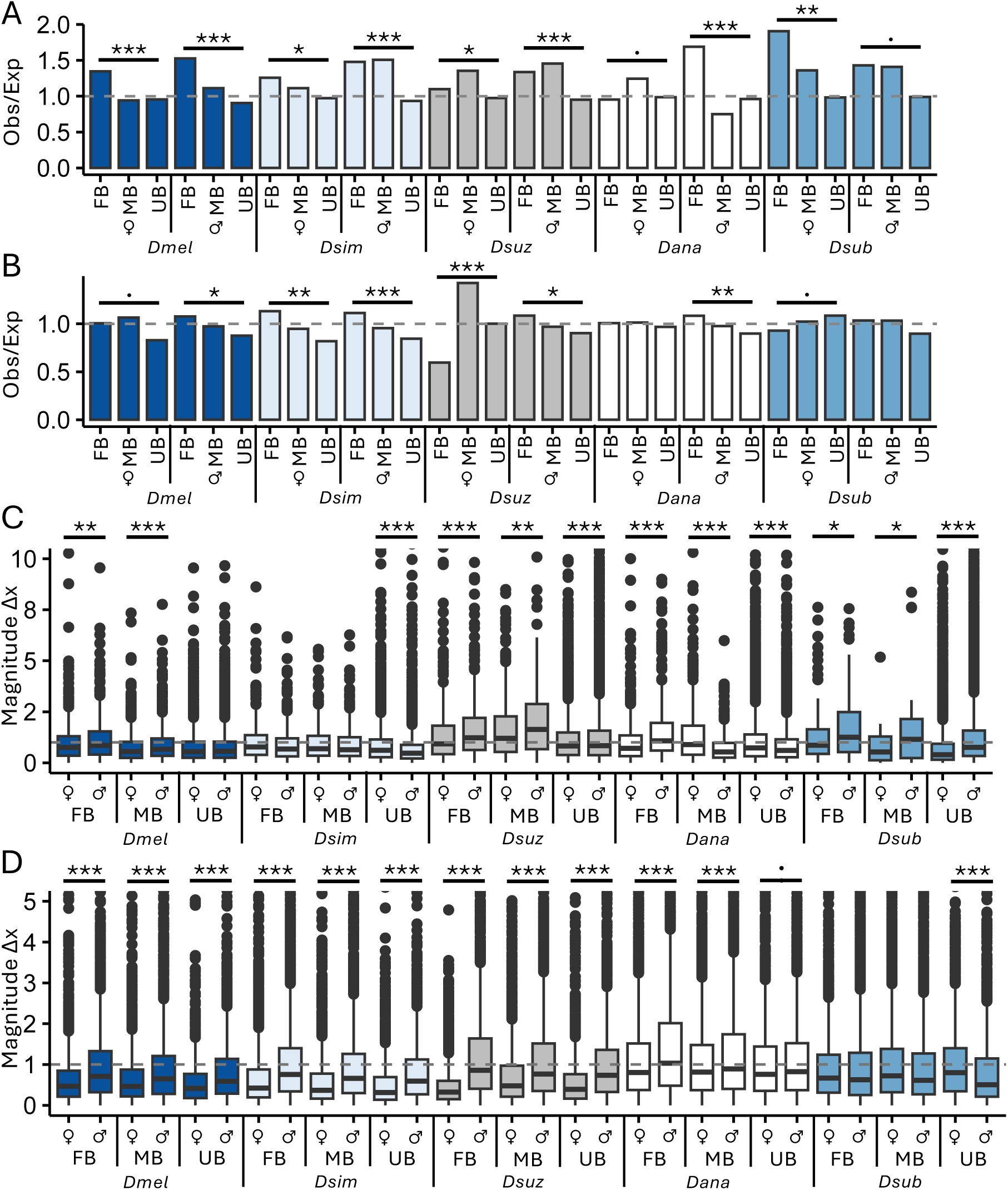
Signs of putative positive selection at the gene expression level. Shown are the ratios of observed (obs) to expected (exp) number of genes under putative positive selection (Δx > 1 or < −1) in each SB category and sex in the A) head and B) body. C, D) Shown is the magnitude of Δx values for FB, MB and UB genes in females and males in the C) head and D) body. The dashed gray line represents the cut-off above which a gene shows signs of putative positive selection (absolute value of Δx > 1). For better visualization, absolute Δx values above 5 are not shown. A, B) Significance was assessed with a χ^2^ test. C, D) Significant differences between sexes were assessed with a *t*-test. \*\*\**P* < 0.005, \*\**P* < 0.01, \**P* < 0.05, marginally non-significant: .*P* < 0.1. Non-significant comparisons not shown.

Next, we examined if signs of putative positive selection could be associated with differences in tissue specificity. We found that increased tissue specificity was associated with an increased magnitude of sex bias, especially in MB genes (Fig 3D). However, a previous study in the brains of 3 *melanogaster* group species found that expression breadth did not selectively constrain the evolution of SB gene expression (Khodursky et al. 2020). To determine if we could detect an association between tissue specificity and evidence of putative positive selection, we compared τ between selected (Δx > 1 or < −1) and non-selected genes in each sex among SB and UB genes in each species and body part. We detected a significant effect of selection regime on tissue specificity (ANOVA, *P* < 0.01 for all; Fig S13). However, in the body, there were no clear overall patterns in the association of tissue specificity and selection regime, except that tissue specificity tended to differ between selected and non-selected genes. In the head, SB genes with signs of putative positive selection tended to show broader expression (i.e. lower τ values), suggesting that the action of selection on more tissue-specific genes may be more constrained than in the body. Thus, although increased tissue specificity was associated with a higher degree of sex bias (Fig 3D), broader expression did not generally preclude the action of positive selection on a SB gene, and was even associated with positive selection in the head (Fig S13).

In order to better understand how turnover in SB expression may be influenced by selection, we examined Δx in genes with lineage-specific gains or losses of sex bias. We detected many of the genes that gained or lost sex bias as under selection for up- or down-regulation, with the exception of SB losses in the head, of which there were few (Table S9). Similar to the results of the full dataset (Fig 6), we often detected similar or higher proportions of genes as under selection in the more lowly expressed sex as in the more highly expressed sex or both sexes (Table S9). However, when we examined if we detected more (or fewer) genes under selection than expected by chance, we only sometimes detected a significant excess of selected genes, which was especially common for male expression during FB gains in the head (χ^2^ test; Table S9). Because we could only include a limited number of genes in this analysis (0–368, mean: 36.9), this general lack of significance is likely at least partially due to a lack of power. Indeed, we could not assess significance for 25–36 of 56 lineage-specific gains/losses due to a lack of power (Tables S9). Because we found that SB expression was often gained via concordant down-regulation in both sexes (Figs 4 and S8), we next examined how often the detected positive selection occurred for a down-regulation of expression in the focal species. We often detected a higher proportion of genes with negative Δx values in 1 or both sexes, but especially in the more lowly expressed sex and especially for male expression of FB genes (Table S9). This finding suggests that selection for sex bias changes may occur fairly frequently for down-regulation, especially in the sex with lower expression. In general, we detected genes under selection for up- or down-regulation in both sexes, suggesting that at least some of the lineage-specific gains and losses that we detected were the result of selection.

## Discussion

We characterized SB gene expression in the head and body of 6 *Drosophila* species, encompassing a broad geographic distribution within species and 2–50 million years of evolution (Fig 1, Table S1). Overall, we detected contrasting patterns of SB gene expression and turnover between the head and body (Table 1, Figs 1–3, S3, S6–S8), as well as differences in expression level, tissue-specificity, and magnitude of sex bias depending on the species, sex bias category, body part and/or sex (Figs S3–S5). Our findings that *i*) sex bias was less extensive (Table 1), *ii*) conserved sex bias was rare while conserved UB expression was extensive (Fig 3A), and *iii*) the least amount of expression divergence occurred in the head (Tables S3–S4) suggest that gene expression is more constrained by shared regulatory mechanisms in the head than in the body. Indeed, sex bias in the head was largely gained in a species-specific manner (Fig 1). Unlike in the body, where sex bias sometimes switched between species, these gains almost exclusively occurred in genes that were either UB or biased towards the same sex in the head of other species (39.2% of examined genes; Fig 3A,B, Figs S6–S7), suggesting that there is greater genetic constraint against sex bias reversals in the head throughout the *Drosophila* genus. Moreover, SB genes in the head were usually also SB in the body irrespective of sex (Fig 3C, Fig S2), suggesting that sex bias is more likely to arise in the head when the underlying regulatory mechanisms for sex bias already exist for that gene. Taken together, these findings suggest that the gain of sex bias is much more genetically constrained in the head than in the body. This pattern is likely driven, at least in part, by expression in the brain, where expression levels are known to be more selectively constrained than in other tissues (Khaitovich et al. 2005, 2006). In keeping with our findings, Khodursky et al. (2020) also found that expression was genetically constrained by shared regulatory mechanisms in the brain with SB expression being largely species-specific; however, they also detected a much larger proportion of switching in sex bias in their dataset. This discrepancy between studies is likely due to their more limited sampling within species, which included either a limited number of isofemale lines (*N* = 2) or lines derived from a single population (*N* = 6). On the other hand, our dataset included more lines (*N* = 5–8) per species, which were derived from multiple, geographically diverse populations. Thus, our samples represented a broader overview of within species variation and likely “averaged out” potential population- or line-specific sex bias. Consistent with this interpretation, a recent study found that somatic SB genes are more variable than UB genes among individuals within sub-species of mice (Xie et al. 2025). This study also detected a larger proportion of sex bias reversals among subspecies, but this discrepancy with our study might be due to their shorter evolutionary timescale (Xi et al. 2025).

Interestingly, lineage-specific gains or losses of sex bias most often occurred via concordant expression changes in both sexes, but with a greater change in 1 sex than the other (75.6% in the head and 54.3% in the body; Fig 4, Figs S9–S10), and this pattern was even stronger for sex bias gains (Fig 4C,F) as well as genes with large expression changes (95.2% in the head and 66.7% in the body; Fig S8), the majority of which occurred via a down-regulation of expression in both sexes. This concordance suggests that these lineage-specific sex bias gains are not a way to resolve sexual antagonism, as in such a case we would expect up-regulated expression in 1 sex and down-regulated expression in the other (quadrants i and iv in Fig 4A,B,D,E). Such changes represented the minority of SB gains in our data (24.4% in the head and 45.7% in the body), especially in the head and among genes with large expression changes (4.8% in the head and 33.3% in the body). Thus, in general, the presence of SB expression is likely to be a poor proxy for ongoing or resolved sexual antagonism in *Drosophila*. In keeping with our findings, a recent study in *Dmel* found that sexually antagonistic effects on starvation resistance could not be explained by SB gene expression (Chen et al. 2025). Moreover, another study of 5 mammalian species found that only 27% of lineage-specific sex bias changes could be associated with turnover in SB transcription factor motifs (Naqvi et al. 2019), suggesting that in such cases the gain of sex-specific regulation may be driving the gain or loss of sex bias. Thus, while opposing sex-specific regulatory changes contribute to the gain or loss of lineage-specific SB gene expression, our results suggest that the majority of lineage-specific SB gains occur via concerted changes in expression with a larger magnitude of change in 1 sex than the other, suggesting that the underlying regulatory architecture is largely shared between the sexes. Indeed, a previous study in humans found that sex differences in the genetic regulation of gene expression were not very common (Oliva et al. 2020), while another study of the brain of 3 *melanogaster* group species found that male and female expression were positively correlated relative to ancestral expression (Khodursky et al. 2020). We similarly detected a positive correlation in lineage-specific SB gains and losses, but the strength of this correlation was stronger for SB gains than losses (Fig 4), underscoring that many species-specific SB gains are likely the result of differences in regulation that is shared between the sexes.

We also detected contrasting patterns in the chromosomal distribution of SB genes between the head and body (Fig 5). In the body, we detected a consistent enrichment of FB and a paucity of MB genes on the X chromosome (Fig 5B), which is consistent with the “feminization” and concurrent “de-masculinization” of the X that has been reported in previous studies (e.g. Parisi et al. 2003; Ranz et al. 2003; Sturgill et al. 2007). On the other hand, in the head, we often detected an enrichment of SB genes (MB, FB, or both) on the X chromosome (Fig 5A). Previous work in *melanogaster* group species found that genes with SB expression, and particularly MB expression, in the brain were enriched on the X chromosome (Catalán et al. 2012; Khodursky et al. 2020) and that this signal from the brain may contribute to a detectable enrichment of X-linked SB genes in studies of whole heads (Huylmans and Parsch 2015). Thus, expression in the brain likely contributed to the enrichment of X-linked SB genes that we detected in the head (Fig 5A).

SB genes, especially MB genes, are often more tissue-specific than UB genes (Meisel 2011; Dean et al. 2016; Khodursky et al. 2020). In keeping with previous findings, we found that tissue specificity was highest in MB genes (Fig S5). Moreover, tissue specificity was positively associated with the magnitude of sex bias, and this pattern was strongest in MB genes, especially in the body (Fig 3D). However, this association did not limit the evolution of sex bias in more broadly expressed genes (Fig S13). Together, these findings suggest that increased tissue specificity may facilitate larger changes in SB expression, especially for MB genes, which is in keeping with the idea that increased tissue specificity may be a driving force in the evolution of SB gene expression, and MB gene expression in particular (Miesel 2011; Dean et al. 2016). However, we should note that in this study we focused on expression in body parts, which are a composite of multiple tissues. While the head is composed of relatively few tissues, the body is composed of many, including reproductive tissues. Moreover, cell composition is more heterogeneous in the body, which together with the tissues it encompasses is composed of more body part-specific cell-type clusters than the head (Li et al. 2022b). Thus, it is possible we may have missed some smaller, tissue-, or cell-type specific changes in SB gene expression. Moreover, this higher heterogeneity may at least in part account for the higher amount of sex bias switching that we detected in the body (14.3% of examined genes versus 0.4% in the head). However, differences in cell-type heterogeneity between the sexes is unlikely to affect our detection of SB genes as a recent study found that cell-type composition differences between the sexes are not a major source of sex bias (Barata and Vicoso 2026).

When we examined Δx as a measure of putative positive selection at the gene expression level (where Δx > 1 or <-1 is indicative of positive selection), we found that *i*) the absolute value of Δx was most often higher in males compared to females, with the mean magnitude of male Δx usually close to or above 1 (Fig 6, Table S8), *ii*) SB genes were often also and sometimes only enriched for signs of positive selection in the opposite sex (i.e. the more lowly expressed sex), especially for male expression of FB genes (Fig 6), and *iii*) Δx tended to shift in the same direction in both sexes (Table S8). At the same time, changes in gene expression relative to ancestral expression during sex bias gains tended to be positively correlated between the sexes (Figs S9–S10), as did gene expression between the sexes within body parts (Tables S3–S4). Taken together, these findings suggest that lineage-specific sex bias changes may often be facilitated by selection acting on expression in both sexes or the sex with lower expression (i.e. selection may be occurring for lower expression, often resulting in a net gain of sex bias in the opposite sex), especially for male expression of FB genes. Indeed, we found that lineage-specific sex bias changes were most often concordant between the sexes, especially in the head, for SB gains, and for genes with large expression changes (Figs 4 and S8), the majority of which occurred via a down-regulation of expression in both sexes. We also detected signs of positive selection in these lineage-specific sex bias changes, especially for male expression of FB genes (Table S9). Thus, our results suggest that most lineage-specific sex bias gains may be the result of selection acting on the asymmetrical down- (or up-) regulation of expression in both sexes via shared regulatory architecture, and this mechanism may be particularly important when expression is highly genetically or selectively constrained, such as in the head or brain, or when there are large overall expression changes. This finding is rather counterintuitive, as SB gene expression is known to evolve rapidly (Assis et al. 2012; Whittle and Johannesson 2013; Harrison et al. 2015; Yang et al. 2016; Tosto et al. 2023; Xie et al. 2025), and theory predicts that shared genetic architecture should slow down the evolution of sexual dimorphism (Lande 1980) given the constraints that the shared genome imposes on gene expression (Griffin et al. 2013; Khodursky et al. 2020). However, a recent study using mouse and human data detected overlapping variances of SB gene expression between the sexes in somatic tissues (Xie et al. 2025), suggesting that patterns and evolutionary dynamics of SB genes are likely more nuanced than previously thought. Therefore, in cases where sex bias is beneficial, but genetic constraint on gene regulation is relatively high, working within the constraint of a shared genome, rather than around it, may provide an alternative, more common route to the establishment of SB gene expression.

## Materials and Methods

### *Drosophila* strains and RNA-seq

All *Drosophila* species were maintained at 21°C with a 14 h light:10 h dark cycle on standard cornmeal–yeast–molasses medium. For all species, strains were established as isofemale lines and maintained by transferring 20–50 flies per generation for at least 10 generations. For all of the *Dmel* and some of the *Dsim* strains, additional inbreeding was done by performing single sib-pair crosses for 5–10 generations. Expression was examined in 5–8 strains of 6 *Drosophila* species (Fig 1). To provide a broad geographical overview of expression within each species, the *Drosophila* strains were sampled from locations across Europe, Asia, and Africa and were either collected by ourselves or kindly provided by others (see Table S1 for details on sampling locations and provenance). Briefly, we included strains from at least 2 populations per species and included strains from the inferred ancestral region for *Dmel*, *Dsim*, *Dana*, and *Dsuz*. For each strain, groups of 30–120 heads and 8–80 bodies of 3–6-day-old flies were dissected in cold 1X PBS and frozen in RNA-DNA shield (Zymo research; Freiburg, Germany) at −70°C until RNA extraction. RNA was extracted using a Qiagen RNeasy Mini or Plus Universal Kit (Qiagen; Hilden, Germany). Poly-A selection, fragmentation, reverse transcription, library construction, and high-throughput RNA-seq was performed by Novogene (Martinsried, Germany) using the Illumina NextSeq 550 platform (Illumina; San Diego, CA) with 150-bp paired-end reads. We sequenced 1 biological replicate per strain, sex, and body part, resulting in 35 libraries for each sample type (140 libraries in total). Library size ranged from 22,874,617 to 34,733,318 million paired-end reads, 43.29–78.74% of which could be mapped, which is relatively low for well-annotated *Drosophila* species and likely due to only mapping to protein coding sequences (CDS) rather than full-length transcripts and ncRNAs (Table S10; see description of read mapping below).

### Read mapping and estimation of gene expression levels

In order to prevent potential bias due to lower quality annotations in some species (for example, no full transcript or ncRNA information was available for *Dimm*; whereas, in *Dmel* UTRs and ncRNAs are better annotated than in the other, less studied species), we mapped RNA-seq reads to CDS sequences (excluding UTRs and ncRNAs) for each species. Reference CDS sequences were downloaded (April 25, 2024) from the following sources: NCBI (*Dsub* genome assembly UCBerk_Dsub_1.0, *Dana* genome assembly ASM1763931v2, D*suz* Genome assembly LBDM_Dsuz_2.1.pri, and *Dsim* genome assembly Prin_Dsim_3.1), FlyBase (version 6.57; Öztürk-Çolak et al. 2024; *Dmel*), and Li et al. (2022a; *Dimm*). Links to the downloaded references are provided in Table S11.

RNA-seq reads were mapped to the respective species reference using NextGenMap (Sedlazeck et al. 2013) in paired-end mode. Read pairs mapping to different genes were discarded, while read pairs matching more than 1 isoform of a gene were randomly assigned to 1 of the isoforms of that gene. For downstream analyses, we analyzed the sum of read counts across all of a gene’s isoforms (across all annotated exons), i.e. on the individual gene-level. In order to make direct comparisons of gene expression among species and sample types, we used our gene counts (Data S1) to calculate levels of gene expression as transcripts per million (TPM; Data S2; Wagner et al. 2013) for all genes in each library, which normalizes for both sequencing depth and potential gene length variation among samples and species. For TPM calculations, CDS lengths were extracted from the *Dmel* reference files or calculated from the reference files for all other species using the faidx function in *samtools* (Danecek et al. 2021). In cases of multiple isoforms, the longest CDS length was used.

### Identification of 1:1 orthologs

In order to keep the datasets for each species comparable, we restricted our analyses to 1:1 orthologs present in all species. To identify 1:1 orthologs, we downloaded (April 29, 2024) pairwise lists of orthologs between *Dmel* and all other species from Li et al. (2022a; for *Dimm*) or DrosOMA (Thiébaut et al. 2024; for all other species; Table S11), as well as the full pairwise orthologs list from DrosOMA (Thiébaut et al. 2024; Table S11). First, all non-1:1 orthologs were removed from each *Dmel* ortholog list and the FlyBase batch download and/or ID validator tools (Öztürk-Çolak et al. 2024) were used to obtain FBgn numbers for *Dmel* IDs from each list. Non-gene or non-coding matches were removed and in cases for which submitted IDs yielded multiple matches, only the FBgn number that exactly corresponded the original ID was retained. For *Dmel* IDs with multiple matches but no exact match or duplicate FBgn numbers due to ID changes/gene re-naming, we used a combination of FlyBase (Öztürk-Çolak et al. 2024), DrosOMA (Thiébaut et al. 2024), and UniProt (Ahmad et al. 2025) to validate the IDs. IDs that did not match between our reference files and our ortholog lists were validated in the same manner. The orthologous *Dmel* sequences corresponding to IDs from the *Dimm* list (Li et al. 2022a) that could not be validated using this method were blasted (blastn; Altschul et al. 1990) to identify the current *Dmel* gene, and their isoform was verified in DrosOMA. All IDs that could not be verified were removed from our 1:1 ortholog list as were IDs that were no longer classified as genes by NCBI, not included in our reference files, and IDs that had been incorporated into a gene model already represented in our list (i.e. duplicate IDs). Next, we used the full pairwise orthologs list from DrosOMA (Thiébaut et al. 2024; Table S11) to remove genes that were not 1:1 orthologs between the non-*melanogaster* species. Orthologs that could not be confirmed as belonging to the same, known OMA group were also removed as were orthologs belonging to the same OMA group (i.e. OMA groups that appeared more than once in our list). After these filtering steps, we obtained a list of 9,016 1:1 orthologs for use in our expression analyses.

### Expression analyses

We identified sex-biased genes using gene counts (Data S1, S3) in the 9,016 1:1 orthologs and a negative binomial test as implemented in DESeq2 (Love et al. 2014) in R (R core team 2022) with a 5% false discovery rate (FDR). In order to exclude lowly expressed genes while retaining as many genes as possible for comparative analyses, genes with low counts or identified as outliers by DESeq2 were removed using the independent filtering function in DESeq2. Because this method retained some lowly expression genes within our analyses (Table S12), we therefore filtered our dataset using a TPM cut-off of **≥** 0.5 for a gene to be considered as expressed and repeated our expression analyses (outlined below). Although applying a TPM cut-off resulted in fewer genes that could be included in the analysis, our results remained qualitatively extremely similar (Tables S12–13 and Fig S14). We therefore focused on our full dataset in the main text and the downstream analyses. To estimate log_2_ fold-changes in gene expression between males and females, head and body samples for each species were analyzed separately using a 1-factor model design with 2 levels corresponding to each sex. For PCAs within each species, we implemented a 2-factor (sex and body part) model design with 2 levels for each factor (male versus female and head versus body). For the PCA including all samples, we similarly implemented a 3-factor design (sex, body part, and species), with the additional third factor consisting of 6 levels corresponding to the 6 examined species. To examine similarities in overall expression between samples (i.e. among body parts, species, and/or sexes), we calculated gene expression divergence between 2 sample types as 1 – Spearman’s ρ between the mean TPM of the 2 sample types across all expressed genes. Significant differences in expression divergence or the absolute value (i.e. magnitude) of sex bias between sample types or sex bias categories were assessed with a *t*-test with a Benjamini-Hochberg correction (Benjamini and Hochberg 1995) for the number of comparisons made per sample or category for each type of data. For each body part, we tested for any effects of sex bias category and/or species on overall expression levels using the *lme4* R package (Bates et al. 2015) and a type II ANOVA as implemented in the Anova function of the *car* (Fox and Weisberg 2019) R package and the log_2_ mean TPM of each gene in each species (log_2_ TPM ∼ species + sex bias category + 1 | gene; Table S13), which provides a more conservative, balanced approach than using TPM values for each individual strain and sex.

To identify lineage-specific gains or losses of sex bias in each body part, we employed a conservative, phylogeny-based approach in which a terminal species (or all species of a terminal clade) was only considered to have a change in sex bias if all species on that lineage shared the same sex bias, and this sex bias differed from that of all other outgroup species, which were also all required to share the same sex bias. Due to the lack of an outgroup, we could not polarize sex bias changes as gains or losses on the lineage leading to *Dimm*; therefore, we categorized the gain or loss of expression relative to the other analyzed species. For example, a gain of SB expression was considered to have occurred in *Dimm* if a gene was SB in *Dimm*, but not in any of the other examined species. For each body part, in order to determine how sex bias changes for individual genes across the entire phylogeny, we calculated the proportion of genes UB, that switched SB anywhere in the phylogeny (i.e were FB and MB in at least 1 species each), and SB in 1-6 of the examined species. Only genes that could be categorized as FB, MB, or UB in all species were included in this analysis, resulting in a total of 7,214 and 8,792 genes in the head and body respectively. To better understand how sex bias changes individually among species, we similarly performed pairwise comparisons between each species using all genes that could be assigned to a sex bias category in both species.

In order to compare how expression changed in a particular species relative to the inferred ancestral expression, we calculated the ancestral expression for each gene, sex and body part for all internal nodes of our phylogenetic tree using the mean TPM values in each species and a Brownian motion model fitted by maximum likelihood as implemented in the ace function in the R package *ape* (Paradis and Schliep 2019). Changes in expression within each sex, species, and body part in comparison to the inferred ancestral expression were calculated as log_2_(TPM_Focal_/TPM_Anc_), where TPM_Focal_ was the mean TPM in the focal species for a given body part and sex and TPM_Anc_ was the inferred ancestral TPM at the nearest internal node. To test for differences in the distribution of expression changes (up- or down-regulation) in males versus females between body parts, we employed a χ^2^ test comparing the number of genes for the entire phylogeny in each quadrant (i–iv, see Fig 4) between the head and body. We further categorized genes as having a large or small expression change by employing a log_2_ fold-change threshold of ≥ 2 in the body or ≥ 1 in the head to categorize a gene as having a large expression change, with values below these thresholds categorized as a small expression change. A lower threshold was applied in the head due to its lower extent (Table 1) and magnitude (Fig S3) of sex bias. To test for significant correlations between male and female expression changes during the gain or loss of sex bias, which would indicate that expression changes are concordant between the sexes, in each body part we calculated Spearman’s ρ between expression changes in males versus changes in females for all genes that gained or lost sex bias within each species. To detect differences in the strength of the association between male and female expression changes when sex bias is gained versus when it is lost, we then tested for differences in strength of these correlations between MB or FB gains and SB losses using a Mann-Whitney *U* test or a paired Mann-Whitney *U* test to test for differences between body parts.

### Chromosomal location and calculation of τ and Δx

Chromosomal locations of all protein-coding genes for *Dsim*, *Dsuz*, *Dana*, and *Dsub* were downloaded (July 11, 2025) from NCBI. For *Dsuz* the CBGP_Dsuzu_IsoJpt1.0 assembly, which is chromosome-level, was used to determine the chromosomal location of analyzed genes. *Dmel* chromosomal locations were extracted from the reference CDS file. Because a chromosome-level assembly was not available for *Dimm*, this species was excluded from all chromosomal location analyses. For all comparisons between X and autosomal genes, we compared genes that could be assigned to the X chromosome (also known as Muller element “A”) to all genes that could be assigned to any autosome, excluding the “dot” chromosome (also known as Muller element “F”), which is thought to have been an ancestral X chromosome (Vicoso and Bachtrog 2013). To detect potential differences in the distribution of SB genes on the X and autosomes for each species, we used a χ^2^ test to test for differences in the number of observed genes in each sex bias category versus the expected number of genes in each category based on the overall proportion of genes in each category for that species separately for the X and the autosomes.

In order to examine tissue specificity, we downloaded the FlyAtlas2 FPKM data (Krause et al. 2022) for all adult *Dmel* tissues, excluding fat body, spermatheca, and heart, which were sequenced using a different chemistry from the other tissues, from https://motif.mvls.gla.ac.uk/downloads/FlyAtlas2_gene_data_2025.xlsx (April 24, 2025). We then calculated the tissue-specificity index τ (Yanai et al. 2005) for each gene (Data S3) and identified the tissue with the highest expression. For each body part, we tested for any effect of sex bias category on τ using a type II ANOVA as implemented in the Anova function of the *car* (Fox and Weisberg 2019) R package (τ ∼ species + sex bias category; Table S13). Because our estimates of τ are the same for every gene regardless of species, we could not include gene or species as a random factor, which would result in overfitting. Moreover, any significant effect of species is a reflection of a different composition of genes in each sex bias category for each species, rather than a direct effect of species on levels of tissue specificity. In order to test for any associations between sex bias and tissue specificity, we calculated Spearman’s ρ between τ and the magnitude of sex bias for each sex bias category in each body part and species. To detect if the sex bias category affects the strength of the association between tissue specificity and the degree of sex bias, we tested for differences in the strength of these correlations among sex bias categories using a Mann-Whitney *U* test.

In order to identify putative signs of positive selection at the gene expression level, for each species and body part we calculated Δx (Data S4) similar to Moghadam et al. (2012) within each sex as

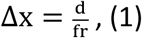

where, *d* is the divergence in gene expression between a focal species and *Dimm*, *r* is the range of expression levels for the focal species, and *f* is a modifier correcting for differences in sample size among species. To calculate Δx, we used the log_2_ TPM as a measure of expression for each gene in each fly strain and calculated the divergence (*d*) in expression between the focal species expression (X_Focal_) and *Dimm* expression (X_Imm_) as

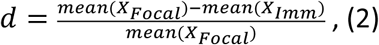

the expression range (*r*) as

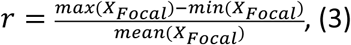

and the correction factor (*f*) as

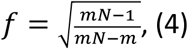

where *m* is the number of strains for the focal species and *N* is the most common number of strains (for our data, *N* = 5). An absolute Δx value above 1 indicates that divergence of expression in the focal species from *Dimm* exceeds expression polymorphism within that species. Thus, a threshold of an absolute Δx value of 1 has previously been taken to indicate that positive selection may have occurred (Moghadam et al. 2012, Zemp et al. 2016, Lichilín et al. 2021, Cossard et al. 2022), with Δx < −1 representing a down-regulation in the focal species and Δx > 1 indicating an up-regulation. To determine if we could detect more genes under putative positive selection than expected by chance, for all genes and for the X chromosome and autosomes separately, we used a χ^2^ test to test for differences in the number of observed putatively positively selected genes in each sex bias category versus the number of expected putatively selected genes based on the proportion of genes in each sex bias category for each sex, species, and body part. To detect potential differences in the overall action of selection in males versus females, for each body part, species, and sex bias category, we used a *t*-test to test for differences in the magnitude of Δx between males and females. To detect potential associations between selection on male versus female expression, we also tested for correlations in Δx between the sexes for each body part, species, and sex bias category using Spearman’s ρ. To determine if body part affects the strength of the association between Δx in each sex, we tested for differences in the strength of these correlations between the body and the head using a paired Mann-Whitney *U* test. To test if putatively selected genes were enriched or depleted among lineage-specific sex bias gains and losses, for expression in each sex we employed a χ^2^ test comparing the observed number of selected and non-selected genes versus the expected number based on the proportion of selected genes in that sample type. Due to lack of power, we excluded cases where the expected number of selected or non-selected genes was below 5 from these analyses. To test if tissue specificity differed between genes identified as under selection versus those identified as not under selection, for each body part and sex, we tested for any effect of selection (i.e. identified as under putative positive selection or not) on τ using a type II ANOVA as implemented in the Anova function of the *car* (Fox and Weisberg 2019) R package (τ ∼ species + sex bias category + selection; Table S13). Because our estimates of τ are the same for every gene regardless of species or sex, we could not include gene or species as a random factor, which would result in overfitting.

## Data Availability

The RNA-seq data that support the findings of this study are publicly available from the European Nucleotide Archive (ENA) at EMBL-EBI under Project accession number XXXXXXXX. All other relevant data are within the manuscript and its Supplementary Material files.

## Acknowledgements

We thank Hilde Lainer for excellent technical assistance in the laboratory. We are grateful to members of the European *Drosophila* Populations Genomics Consortium (DrosEU) and the following people for providing the fly lines used in this study: Jessica Abbott, Nicolas Gompel, Josefa González, Sonja Grath, Katja Hoedjes, Mihailo Jelić, Maaria Kankare, Viola Nolte, Darren Obbard, Marta Pascual, John Pool, Christian Schlötterer, Mads Fristrup Schou, Marina Stamenković, Fabian Staubach, Cristina Vieira, Jorge Vieira, and Ricardo Wilches. This study was supported by the Deutsche Forschungsgemeinschaft priority program “The genomic basis of evolutionary innovations” (SPP2349; Project No. 503272152 awarded to JP).

## Supplementary Material Captions

**Table S1:** Overview of *Drosophila* lines assayed in this study

**Table S2:** Sex biased genes in *D. melanogaster* in a random sub-sampling of 5 lines

**Table S3:** Expression divergence within species

**Table S4:** Expression divergence among species

**Table S5:** Number of SB genes with highest expression in testis, ovary, or a somatic tissue

**Table S6:** Lineage-specific gains and losses of sex biased gene expression

**Table S7:** Sex-biased genes on the X versus autosomes

**Table S8:** Genes showing signs of putative positive selection at the expression level

**Table S9:** Putative positive selection in genes with lineage-specific SB gains and losses

**Table S10:** Library size and mapping efficiency

**Table S11:** Download location of files used for analysis

**Table S12:** Sex bias in the examined species using TPM cut-off

**Table S13:** Results of Type II ANOVAs to test the effect of sex bias category and species on gene expression levels and τ

**Fig S1:** Principal component analysis of gene expression profiles in A) *Dmel*, B) *Dsim*, C) *Dsuz*, D) *Dana*, E) *Dsub*, and F) *Dimm*. Triangles and diamonds indicate female and male body, respectively, while circles and squares indicate female and male head, respectively.

**Fig S2:** Overlap in SB genes between head (H) and body (B). Shown are upset plots demonstrating the overlap in UB, FB, or MB genes in A) *Dmel* (mel), B) *Dsim* (sim), C) *Dsuz* (suz), D) *Dana* (ana), E) *Dsub* (sub), and F) *Dimm* (imm). Horizontal bars represent the total number of genes in a body part and sex bias category combination. Vertical bars represent the number of genes in an intersection class. Connected, filled circles underneath a vertical bar indicate that a body part and sex bias category combination is included in an intersection class. Blue represents genes UB in one body part and SB in the other, dark gray represents gene in the same bias category, and light blue indicates genes with the opposite sex bias between head and body.

**Fig S3:** Magnitude of sex bias in the A) head and B) body of *D. melanogaster* (mel), *Dsim* (sim), *Dsuz* (suz), *Dana* (ana), *Dsub* (sub), and *Dimm* (im/imm). Significant differences between species for MB and FB genes were assessed with a *t*-test. Shown are the BH-corrected *P*-values in the C) head and D) body. Non-significant *P*-values are shown in bold.

**Fig S4:** Overall expression levels in the A) head and B) body for MB (M), FB (F), and UB (U) genes in *Dmel*, *Dsim*, *Dsuz*, *Dana*, *Dsub*, and *Dimm*. Significance was assessed for each body part with a type II ANOVA with sex bias (SB) category and species as fixed factors and gene as a random factor.

**Fig S5:** Tissue specificity τ in the A) head and B) body for MB (M), FB (F), and UB (U) genes in *Dmel*, *Dsim*, *Dsuz*, *Dana*, *Dsub*, and *Dimm*. Significance was assessed for each body part with a type II ANOVA with sex bias (SB) category and species as factors.

**Fig S6:** Overlap in SB genes within the head (H) between species. Shown are upset plots demonstrating the overlap in UB, FB, or MB genes in pairwise comparisons between A–E) *Dmel* (mel), A, F–I) *Dsim* (sim), B,F,J–L) *Dsuz* (suz), C,G,J,M,N) *Dana* (ana), D,H,K,M,O) *Dsub* (sub), and E,I,L,N,O) *Dimm* (imm). Horizontal bars represent the total number of genes in a species and sex bias category combination. Vertical bars represent the number of genes in an intersection class. Connected, filled circles underneath a vertical bar indicate that a species and sex bias category combination is included in an intersection class. Blue represents genes UB in one species and SB in the other, dark gray represents genes in the same sex bias category, and light blue indicates genes that switch sex bias between species. P) Summary of pairwise overlap in sex bias categories among species. Shown are the proportion of overlapping genes that had the same SB (white) or were both UB (dark blue), were SB in only one of the examined species (SB 1 species; light blue), or switched sex bias (Switch SB; gray) between *Dmel* (*m*), *Dsim* (*si*), *Dsuz* (*su*), *Dana* (*a*), *Dsub* (*sb*), and *Dimm* (*im*).

**Fig S7:** Overlap in sex biased genes within the body (B) between species. Shown are upset plots demonstrating the overlap in UB, FB, or MB genes in pairwise comparisons between A– E) *Dmel* (mel), A, F–I) *Dsim* (sim), B,F,J–L) *D. suz* (suz), C,G,J,M,N) *Dana* (ana), D,H,K,M,O) *Dsub* (sub), and E,I,L,N,O) *Dimm* (imm). Horizontal bars represent the total number of genes in a species and sex bias category combination. Vertical bars represent the number of genes in an intersection class. Connected, filled circles underneath a vertical bar indicate that a species and sex bias category combination is included in an intersection class. Blue represents genes UB in one species and SB in the other, dark gray represents genes in the same sex bias category, and light blue indicates genes that switch sex bias between species. P) Summary of pairwise overlap in sex bias categories among species. Shown are the proportion of overlapping genes that had the same SB (white) or were both UB (dark blue), were SB in only one of the examined species (SB 1 species; light blue), or switched sex bias (Switch SB; gray) between *Dmel* (*m*), *Dsim* (*si*), *Dsuz* (*su*), *Dana* (*a*), *Dsub* (*sb*), and *Dimm* (*im*).

**Fig S8:** All SB gains and losses for genes with A,B) large and C,D) small expression changes between the focal and ancestral species in the A,C) head and B,D) body. The cut-off for a large expression change was a log_2_ fold-change of magnitude 1 or higher in the head and 2 or higher in the body. Genes with concordant (con) changes in expression between the sexes are located in quadrants ii and iii; while genes with opposing (opp) expression changes between the sexes are located in quadrants i and iv.

**Fig S9:** Expression changes during sex bias turnover in the head. Shown are scatterplots of expression in A–C) *Dmel*, D–F) *Dsim*, G–I) *Dsuz*, J–L) *Dana*, M–O) *Dsub*, and P–R) *Dimm* in each sex relative to ancestral (anc) expression for A,D,G,J,M,P) FB or B,E,H,K,N,O) MB gains or C,F,I, L,O,R) losses. Spearman’s ρ and the associated *P* value for each category are shown.

**Fig S10:** Expression changes during sex bias turnover in the body. Shown are scatterplots of expression in A–C) *Dmel*, D–F) *Dsim*, G–I) *Dsuz*, J–L) *Dana*, M–O) *Dsub*, and P–R) *Dimm* in each sex relative to ancestral (anc) expression for A,D,G,J,M,P) FB or B,E,H,K,N,O) MB gains or C,F,I, L,O,R) losses. Spearman’s ρ and the associated *P* value for each category are shown.

**Fig S11:** Δx in the A) head and B) body of 5 *Drosophila* species. Shown are Δx values for FB, MB and UB genes in females (F) and males (M) of *Dmel*, *Dsim*, *Dsuz*, *Dana*, and *Dsub*. The dashed gray line represents the cut-off above or below which a gene shows signs of putative positive selection. For better visualization, Δx values above 5 or below −5 are not shown.

**Fig S12:** Chromosomal distribution of signs of putative positive selection at the gene expression level. Shown are the ratios of observed (obs) to expected (exp) number of genes under putative positive selection (Δx > 1 or < −1) in each sex bias category and sex on the X chromosome and autosomes (auto) in the A) head and B) body. Significance was assessed with a χ^2^ test. \*\*\**P* < 0.005, \*\**P* < 0.01, \**P* < 0.05, marginally non-significant: .*P* < 0.1. Non-significant comparisons not shown.

**Fig S13:** Tissue specificity, τ, in genes identified as putatively under positive selection (S) or not under selection (N) in A,C) females (F) and B,D) males (M) the A,B) head and C,D) body for MB (M), FB (F), and UB (U) genes in *Dmel*, *Dsim*, *Dsuz*, *Dana*, *Dsub*, and *Dimm*. Significance was assessed for each body part with a type II ANOVA with sex bias (SB) category, species, and if under putative positive selection as factors.

**Fig S14:** Expression analyses using a TPM cut-off of 0.5. A–D) Shown are A,B) overall expression levels and C,D) tissue specificity, τ, in the A,C) head and B,D) body for MB (M), FB (F), and UB (U) genes in *Dmel* (*mel*), *Dsim* (*sim*), *Dsuz* (*suz*), *Dana* (*ana*), *Dsub* (*sub*), and *Dimm* (*imm*). A,B) Significance was assessed for each body part with a type II ANOVA with sex bias (SB) category and species as fixed factors and gene as a random factor. C,D) Significance was assessed for each body part with a type II ANOVA with SB category and species as factors. E,F) Shown are Spearman’s ρ correlations between male and female expression changes within each species for SB gains and losses in the E) head (H) and F) body (B). Significance was assessed with a Mann-Whitney U test. ***P < 0.005, **P < 0.01, *P < 0.05. Non-significant comparisons not shown. G–L) Genes with concordant (con) changes in expression between the sexes are located in quadrants ii and iii; while genes with opposing (opp) expression changes between the sexes are located in quadrants i and iv. Total genes in each quadrant for the entire phylogeny in the G) head and H) body are shown. In parentheses, the proportion of total gene expression changes are shown. G–J) Shown are all SB gains and losses for genes with I,J) large and K,L) small expression changes between the focal and ancestral species in the I,K) head and J,L) body. The cut-off for a large expression change was a log_2_ fold-change of magnitude 1 or higher in the head and 2 or higher in the body.

**Data S1:** Data underlying gene expression analyses: gene counts.

**Data S2:** Data underlying gene expression analyses: TPM.

**Data S3:** Data underlying gene expression analyses: SB gene expression, τ, and expression changes.

**Data S4:** Data underlying selection analyses.

